# Clinical metagenomic identification of *Balamuthia mandrillaris* encephalitis and assembly of the draft genome: the critical need for reference strain sequencing

**DOI:** 10.1101/024455

**Authors:** Alexander L. Greninger, Kevin Messacar, Thelma Dunnebacke, Samia N. Naccache, Scot Federman, Jerome Bouquet, David Mirsky, Yosuke Nomura, Shigeo Yagi, Carol Glaser, Michael Vollmer, Craig A. Press, Bette K. Klenschmidt-DeMasters, Samuel R. Dominguez, Charles Y. Chiu

## Abstract

Primary amoebic meningoencephalitis (PAM) is a rare, often lethal cause of encephalitis, for which early diagnosis and prompt initiation of combination antimicrobials may improve clinical outcomes. In this study, we present the first draft assembly of the *Balamuthia mandrillaris* genome recovered from a rare survivor of PAM, in total comprising 49 Mb of sequence. Comparative analysis of the mitochondrial genome and high-copy number genes from 6 additional *Balamuthia mandrillaris* strains demonstrated remarkable sequence variation, with the closest homologs corresponding to other amoebae, hydroids, algae, slime molds, and peat moss,. We also describe the use of unbiased metagenomic next-generation sequencing (NGS) and SURPI bioinformatics analysis to diagnose an ultimately fatal case of *Balamuthia mandrillaris* encephalitis in a 15-year old girl. Real-time NGS testing of a hospital day 6 CSF sample detected *Balamuthia* on the basis of high-quality hits to 16S and 18S ribosomal RNA sequences present in the National Center for Biotechnology Information (NCBI) nt reference database. Retrospective analysis of a day 1 CSF sample revealed that more timely identification of *Balamuthia* by metagenomic NGS, potentially resulting in a better outcome, would have required availability of the complete genome sequence. These results underscore the diverse evolutionary origins underpinning this eukaryotic pathogen, and the critical importance of whole-genome reference sequences for microbial detection by NGS.

## BACKGROUND

*Balamuthia mandrillaris* is a free-living amoeba that is a rare, almost uniformly lethal, cause of primary amoebic encephalitis (PAM) in humans (Visvesvara 1993). Originally isolated from the brain of a baboon at the San Diego Zoo in 1986, *Balamuthia mandrillaris* has since been reported in over 100 cases of PAM worldwide (Visvesvara et al. 1990; Schuster et al. 2006; Matin et al. 2008), and amoebae associated with fatal encephalitis in a child have been cultured directly from soil (Schuster et al. 2003). The vast majority of cases are fatal, although there are a few published case reports of patients surviving *Balamuthia* encephalitis after receiving combination antimicrobial therapy and in vitro data supporting the potential efficacy of several antimicrobial agents (Schuster and Visvesvara 1996; Deetz et al. 2003; Schuster et al. 2006; Martinez et al. 2010; Ahmad et al. 2013; Kato et al. 2013). Despite the availability of validated RT-PCR assays for the detection of free-living amoebae (Yagi et al. 2005; Qvarnstrom et al. 2006), PAM is not often clinically suspected and the diagnosis is most commonly made around the time of death or post-mortem on brain biopsy (Schuster et al. 2004; Perez and Bush 2007).

Our lab has demonstrated the capacity of metagenomic next-generation sequencing (NGS) to provide clinically actionable information in a number of acute infectious diseases, most notably encephalitis (Wilson et al. 2014; Naccache et al. 2015). This approach enables the rapid and simultaneous detection of viruses, bacteria, and eukaryotic parasites in clinical samples (Naccache et al. 2014). Encephalitis is a critical illness with a broad differential, for which unbiased diagnostic tools such as metagenomic NGS can make a significant impact (Schubert and Wilson 2015). However, the utility of diagnostic NGS is highly dependent on the breadth and quality of databases that contain whole-genome sequence information of reference strains needed for alignment (Fricke and Rasko 2014).

In this study, we describe the first draft genome sequence of a strain of *Balamuthia mandrillaris* from a rare survivor of PAM and comparative sequence analysis of 6 additional mitochondrial genomes. We also demonstrate the ability of metagenomic NGS to rapidly detect *Balamuthia mandrillaris* from the cerebrospinal fluid (CSF) of an critically ill 15-year old, and highlight the importance of genomic reference sequences by retrospective analysis of a hospital day (HD) 1 sample.

## RESULTS

### First mitochondrial genome of *Balamuthia mandrillaris*

We cultured *Balamuthia mandrillaris* strain 2046 in axenic media from a brain biopsy corresponding to a rare survivor of PAM (Vollmer and Glaser, manuscript in review). Sequencing of DNA extracted from the axenic culture generated 3.8 million 75 base pair (bp) mate-pair reads with an average insert size of 2,187 bp. *De novo* assembly yielded a circular mitochondrial genome of 41,656 bases that was comprised of 64.8% AT at 2,082X coverage (Fig. 1A). The overall size and AT content of the *Balamuthia mandrillaris* mitochondrial genome was closer to that of *Acanthamoeba castellani* (41,591 bp, 70.6% AT) (Burger et al. 1995) than *Naegleria fowleri* (49,531 bp, 74.8% AT) (Herman et al. 2013), although overall average nucleotide identity with *Balamuthia* was found to be low for both amoebae (∼68%).

**Figure 1.**
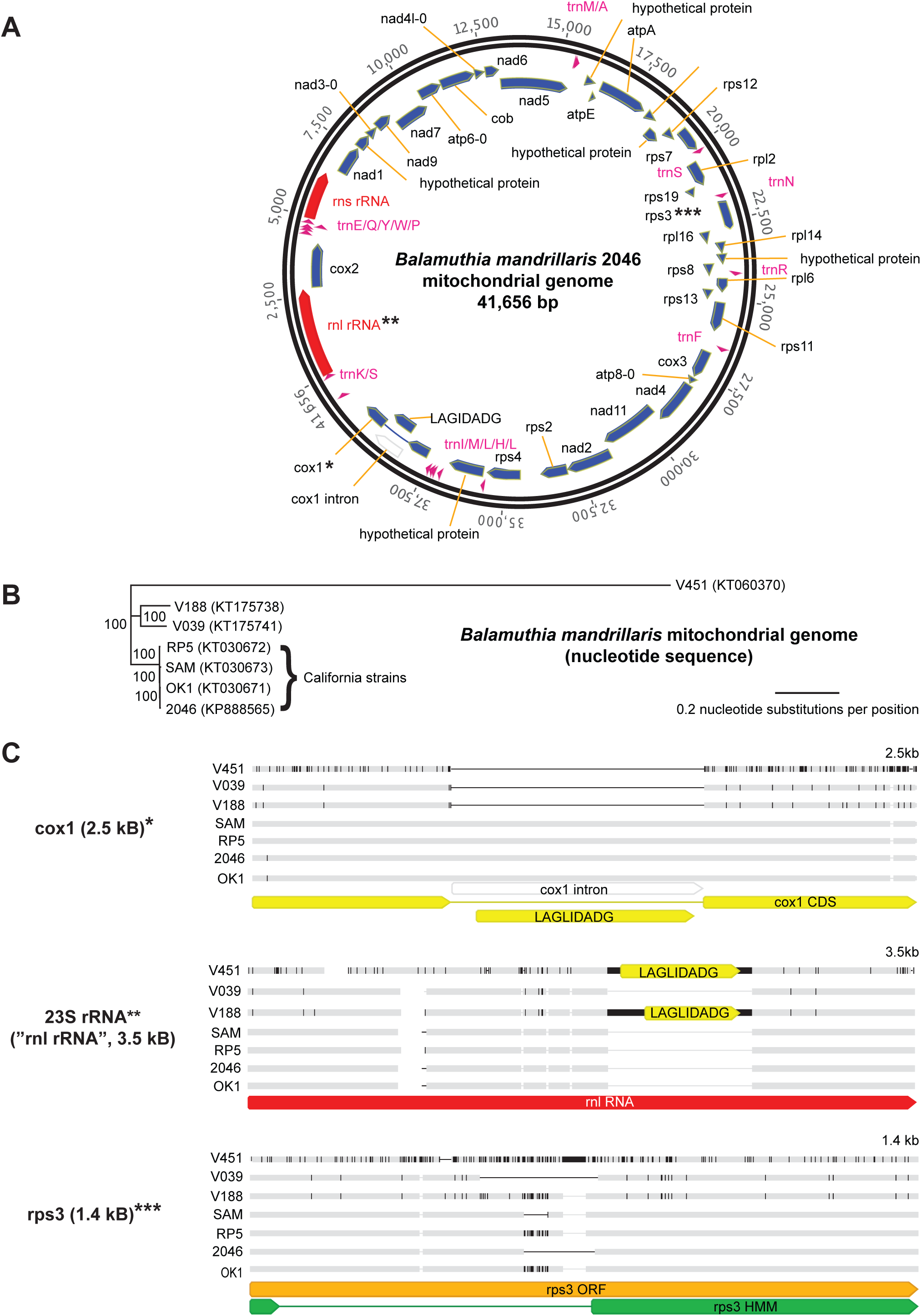
Sequencing and comparative phylogenetic analysis of the mitochondrial genome of *Balamuthia mandrillaris*. **(A)** *Balamuthia mandrillaris* 2046 mitochondrial genome. Annotation of the 41,656 bp genome was performed using RNAmmer, tRNAscan-SE, and Glimmer gene predictor, with all ORFs manually verified using BLASTx alignment. (**B)** Phylogenetic analysis of 7 newly sequenced genomes from different strains of *Balamuthia mandrillaris*. An outgroup (e.g. *Acanthamoeba castellani*) is not shown given the lack of gene synteny. Branch lengths are drawn proportionally to the number of nucleotide substitutions per position, and support values are shown for each node. **(C)** Differences in individual gene features (cox1, 23S rRNA, and rps3), among the 7 mitochondrial genomes, as detailed in the text.

The *Balamuthia mandrillaris* 2046 mitochondrial genome contained 2 ribosomal RNA (rRNA) genes, 18 transfer RNA (tRNA) genes, and 38 coding sequences, with 5 of those being hypothetical proteins. The organization of the mitochondrial genome retained several syntenic blocks with the *Acanthamoeba castellani* genome, including tnad3-9-7-atp6 and rpl11-rps12-rps7-rpl2-rps19-rps3-rpl16-rpl14. However, many other features of the genome were unique, such as the order of the remaining coding blocks, the lack of a combined cox1/cox2 gene, as present in *Acanthamoeba castellani* (Burger et al. 1995), and the lack of intron splicing in 23S rRNA. The *Balamuthia mandrillaris* mitochondrial cox1 gene was interrupted by a LAGLIDADG endonuclease open reading frame (ORF) containing a group I intron, as has been reported for a wide variety of other eukaryotic species (Fukami et al. 2007; Zheng et al. 2012). The putative rps3 gene was encoded within a 1290 bp ORF that, when translated, aligned by hidden Markov model (HMM) analysis to rps3 proteins from *Escherichia coli* (PDB 4TP8/4U26) and *Thermus thermophilus* (PDB 4RB5/4W2F), and only in base positions 1-66 and 583-1290 of the ORF. This finding was consistent with the presence of a putative >500 bp intron in the *Balamuthia mandrillaris* rp3 gene that to date has only been described to in plants (Laroche and Bousquet 1999; Regina et al. 2005). Alternatively, the ORF was also found to encode a putative tRNA (Asn) such that the 5’ end of the ORF could represent a large intergenic sequence.

### Mitochondrial Genome Diversity of Balamuthia mandrillaris

To investigate the extent of sequence diversity in *Balamuthia mandrillaris*, we sequenced the mitochondrial genomes from 6 additional *Balamuthia* strains available at the California Department of Public Health and the American Tissue Culture Collection (Table 1). The 7 total circular mitochondrial genomes averaged 41,526 bp in size (range 39,996 to 42,217 bp), and shared pairwise nucleotide identities ranging from 82.6% to 99.8%. The phylogeny revealed the presence of at least 3 separate lineages of *Balmuthia mandrillaris*, with all of the strains from California that had been submitted to the California Department of Public Health clustering together in a single clade (Fig. 1B). Consistent with a previous report (Booton et al. 2003), we found that the mitochondrial genome of strain V451 was the most divergent among tested strains, and possessed an additional 1,149 bp ORF downstream of the cox1 gene that did not align significantly to any sequence in the NCBI nt or nr reference database.

**Table 1.** Strains used in this study.

| Strain | Date | Location | Note | Citation |
| --- | --- | --- | --- | --- |
| SAM** | Spring 2001 | Rohnert Park, CA | 3 yo girl, cultured on Vero cells | Bakardjiev A et al. 2002 |
| RP-5** | Spring 2001 | Rohnert Park, CA | environmental sample associated with SAM, cultured on Vero cells | Schuster FL et al. 2003 |
| OK1** | 2002 | Oakland, CA | environmental sample in NorCal unrelated to SAM, cultured on Vero cells | Dunnebacke TH et al. 2004 |
| V039* | 1990 | San Diego, CA | type strain isolated from pregnant mandrill at San Diego Zoo, cultured on Vero cells | Visvesvara GS et al. 1990 |
| V188** | 1989 | Georgia | 59 yo man, amoeba isolated from brain/skin lesion, cultured on Vero cells | Gordon SM et al. 1992 |
| V451** | N/A | New York | 6 yo girl, cultured on Vero cells | Yang XH et al. 2001 |
| 2046** | March 2010 | Walnut Creek, CA | 26 yo man, survivor, cultured on Vero cells | Vollmer and Glaser, in review |
| CSF108*** | May 2015 | Colorado | 15 yo girl, direct metagenomic detection from CSF |  |

\*strain obtained from ATCC (V39)
\*\*strain obtained from California Department of Public Health
\*\*\*strain obtained from recent clinical case in Colorado

Putative introns constituted the major source of variation among the mitochondrial genomes (Fig. 1C). Four strains out of 7, including strain 2046 from the rare survivor of *Balamuthia* infection, contained a 975 bp LAGLIDADG-containing intron in the cox1 gene, whereas no such intron was present in the remaining 3 strains. Two of the remaining 3 strains, strains V451 and V188, instead had an approximately 790 bp insert in the 23S rRNA gene (Figure 1A, “rnl RNA”) that contained a 530 bp or 666 bp LAGLIDADG-containing ORF, respectively, and that coded for a putative endonuclease. The LAGLIGDADG-containing endonucleases in the 2 strains shared 84% amino acid pairwise identity with each other, but ∼50% amino acid identity to a corresponding LAGLIGDADG-containing endonuclease in the 23S rRNA gene of *Acanthamoeba castellani*, and <12% amino acid identity to the LADGLIDADG-containing cox1 introns in the four other *Balamuthia* strains. The final remaining strain, ATCC-V039, lacked an intron in either the cox1 or 23S rRNA gene.

The ORF containing the rps3 gene, found to contain a possible rps3 intron or intergenic region by analysis of the strain 2046 mitochondrial genome, varied in length among the 7 sequenced mitochondrial genomes from 1,290 bp to 1,425 bp. Notably, the length of the putative intron or intergenic region accounted for all of the differences in overall length of the rps3 gene. Confirmatory PCR and sequencing of this locus using conserved outside primers revealed that each strain tested had a unique length and sequence (Fig. 2), raising the possibility of targeting this region for *Balamuthia* mandrillaris strain detection and genotyping.

**Figure 2.**
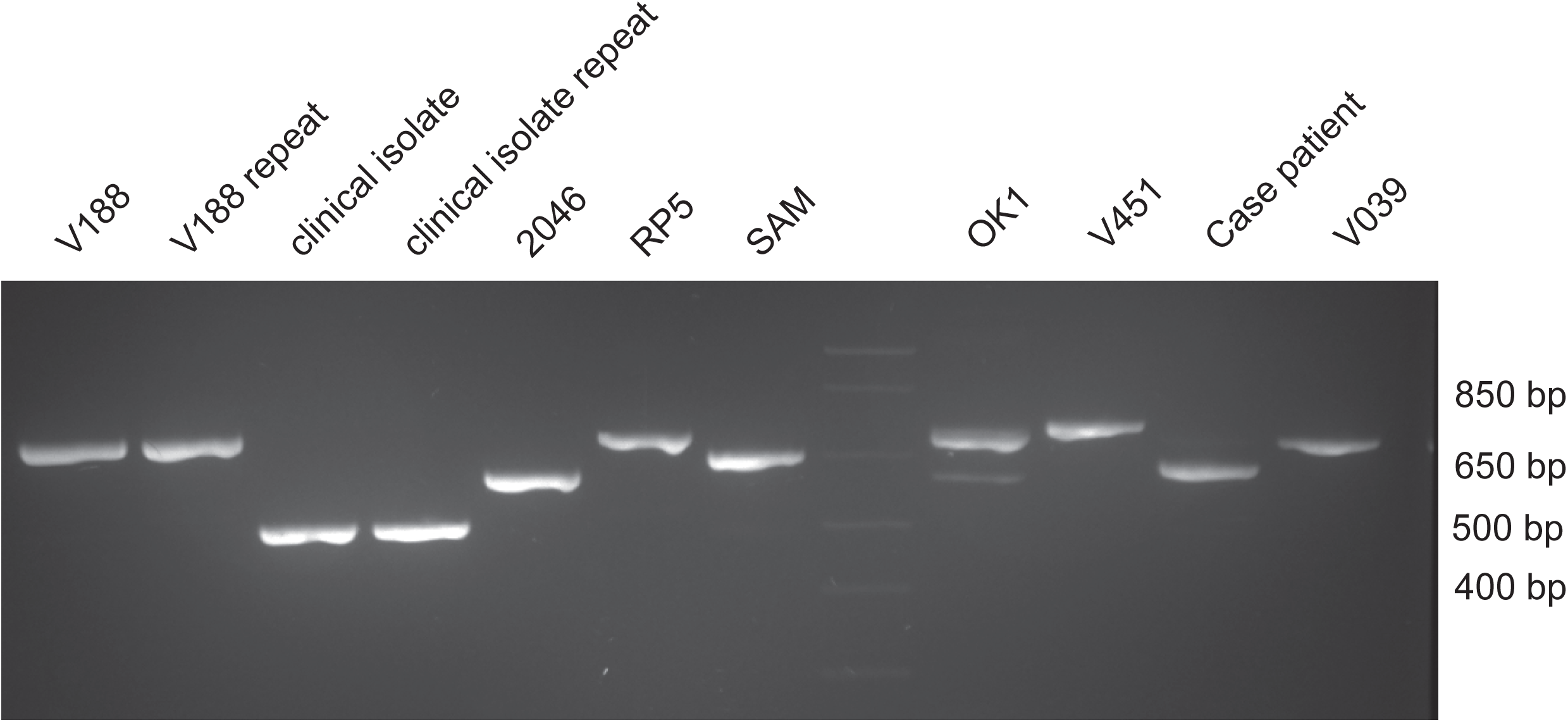
PCR amplification of the *Balamuthia* rps3 mitochondrial gene. The variable length of the rps3 intron among 8 different *Balamuthia* strains (7 newly sequenced mitochondrial genomes and the case patient) suggests that this gene may be an attractive target for development of a molecular genotyping assay. Column 4 corresponds to the DNA ladder (faint appearance), while columns 2 and 3 correspond to an additional clinical *Balamuthia* isolate whose mitochondrial genome was not sequenced.

### First draft genome of *Balamuthia mandrillaris*

Because of the high-copy number of mitochondrial sequences in *Balamuthia*, as noted previously for *Naegleria fowleri* (Herman et al. 2013), we performed an additional NGS run of 14.1 million 250 bp single-end reads, and computationally subtracted reads aligning to the mitochondrial genome. Assembly of the remaining 4.4 million high-quality reads yielded 31,194 contiguous sequences (contigs) with an N50 of 3,411 bp. Scaffolding and gap closure using an additional 57.4 million NGS reads and computational removal of exogenous sequence contaminants yielded a final assembly of ∼44.3 Mb comprised of 14,699 scaffolds with an N50 of 19,012 bp (Table 2). Direct BLASTn alignment of the scaffolds to the National Center for Biotechnology Information (NCBI) nt database revealed that the most common organism aligning to *Balamuthia mandrillaris* was *Mus musculus* (house mouse) (2,067/14,699 = 14.1% of scaffolds), nearly entirely due to low-complexity sequences, followed by high-significance hits to *Acanthamoeba castellani* (627/14,699 = 4.3% of scaffolds).

**Table 2.** Sequencing runs and genome assembly details.

| Sequencing run | Raw reads | Filtered reads | Read Length | Library Prep |
| --- | --- | --- | --- | --- |
| MP1* | 3845734 | N/A | 2x80bp | Nextera Mate-Pair |
| MP2* | 31223454 | 12095792 | 2x80bp | Nextera Mate-Pair |
| MP3* | 14121900 | 4397510 | 2x250bp | Nextera Mate-Pair |
| PE1** | 213604902 | 29314946 | 2x250bp | Nextera XT |
| Assembly*** | Contigs | N50 (bp) |  |  |
| Mitochondria | 1 | 41,656 | 41,656 |  |
| contigs | 31,194 | 3,411 | 48938887 |  |
| scaffolds | 26,811 | 19,415 | 49120517 |  |
| gap-fill | 26,811 | 19,012 | 48939625 |  |
| Final assembly (minus <500bp, contaminants) | 14,699 | 26,144 | 44270879 |  |

\*3 runs of a Nextera mate-pair library of strain 2046 from axenic culture sequenced on an Illumina MiSeq
\*\*Nextera XT paired-end library of strain 2046 sequenced on an Illumina HiSeq
\*\*\*assembly of mitochondrial genome from MP1 library, of whole-genome from all 4 libraries

Analysis of individual genes from the *Balamuthia* mitochondrial genome revealed the presence of significant diversity across all kingdoms of life (Fig. 3). The 18S-28S rRNA locus in the *Balamuthia* mitochondrial genome corresponded to a 12.5 kB contig sequenced at high coverage (>400X over rRNA regions). The previously sequenced 18S gene (2,017 bp) demonstrated 99.5% identity to existing *Balamuthia mandrillaris* 18S rRNA sequences in the NCBI nt database, while the 28S gene (4999 bp) had homology across multiple diverse species, with only 68.5% pairwise identity to its closest phylogenetic relative, *Acanthamoeba castellani* (Fig. 3B). From the nuclear genome, one high-copy contig contained a truncated 5,250 nucleotide ORF exhibiting only 33% amino acid identity to *Rhizopus delemar* (pin mold), and harboring elements consistent with a retrotransposon (Kordis 2005), including an RNAse HI from Ty3/Gypsy family retroelements, a reverse transcriptase, a chromodomain, and a retropepsin. Two high-copy, ∼1,600 bp ORFs that failed to match any sequence by BLASTx alignment to the NCBI nr protein database were found to align significantly to *Escherichia coli* site-specific recombinase by remote homology HMM analysis.

**Figure 3.**
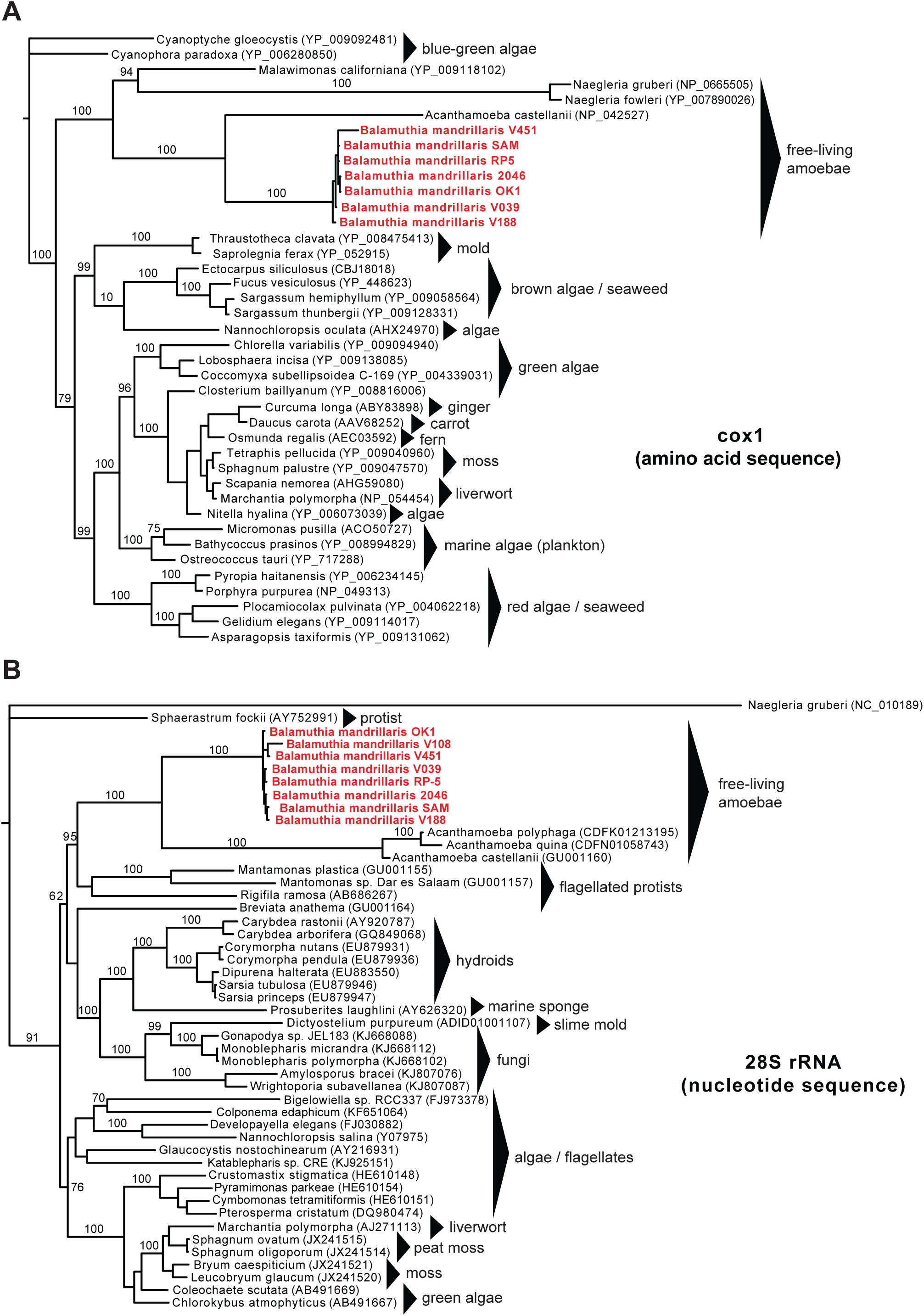
Phylogenetic trees of the mitochondrial cox1 protein and 28S rRNA gene reveal the close phylogenetic relationship between *Balamuthia* and *Acanthamoeba*. **(A)** Phylogeny of 7 *Balamuthia* cox1 amino acid sequences along with the top complete sequence hits in NCBI nr ranked by BLASTp E-score. **(B)** Phylogeny of 7 *Balamuthia* 23S rRNA nucleotide sequences along with the top complete sequence hits in NCBI nt ranked by BLASTn E-score. Sequences were aligned using MUSCLE and a phylogenetic tree constructed using MrBayes. Branch lengths are drawn proportionally to the number of nucleotide substitutions per position, and support values are shown for each node.

### A case of *Balamuthia* encephalitis diagnosed by metagenomic NGS

Concurrent with assembly of the *Balamuthia* genome, metagenomic NGS was performed to investigate a case of meningoencephalitis in a 15-year-old girl with insulin-dependent diabetes mellitus and celiac disease. The patient initially presented to a community emergency room with 7 days of progressive symptoms including right arm weakness, headache, vomiting, ataxia, and confusion. Her diabetes was well controlled with an insulin pump, and she did not take any additional medications. Exposure history was significant for contact with alpacas at a family farm and swimming in a freshwater pond nine months prior. She had no international travel, sick contacts, or insect bites. She was given 10 mg dexamethasone with symptomatic improvement in her headaches, but was subsequently transferred to a tertiary care children’s hospital after a computed tomography scan revealed left occipital and frontal hypodensities.

On HD 1, peripheral white blood count was 11.6x10^3^ cells/μL (89% neutrophils, 6% lymphocytes, 4% monocytes), erythrocyte sedimendation rate was 13 mm/hr [normal range 0-20 mm/hr], C-reactive reactive was 3 mg/dL [normal range 0-1 mg/dL], and procalcitonin 0.05 ng/mL [normal range 0-0.5 ng/mL]. CSF analysis demonstrated 377 leukocytes/μL (2% neutrophils, 53% lymphocytes, 39% monocytes, and 6% eosinophils), glucose of 122 mg/dL [normal range, 40-75 mg/dL], and protein of 59 mg/dL [normal range, 12-60 mg/dL]. Viral polymerase chain reaction (PCR) testing for herpes simplex virus (HSV) and bacterial cultures were negative. Magnetic resonance imaging (MRI) scan of the brain on HD 1 showed hemorrhagic lesions with surrounding edema in the superior left frontal lobe and left occipital lobe with a small focus of edema in the right cerebellum (Fig. 4A).

**Figure 4.**
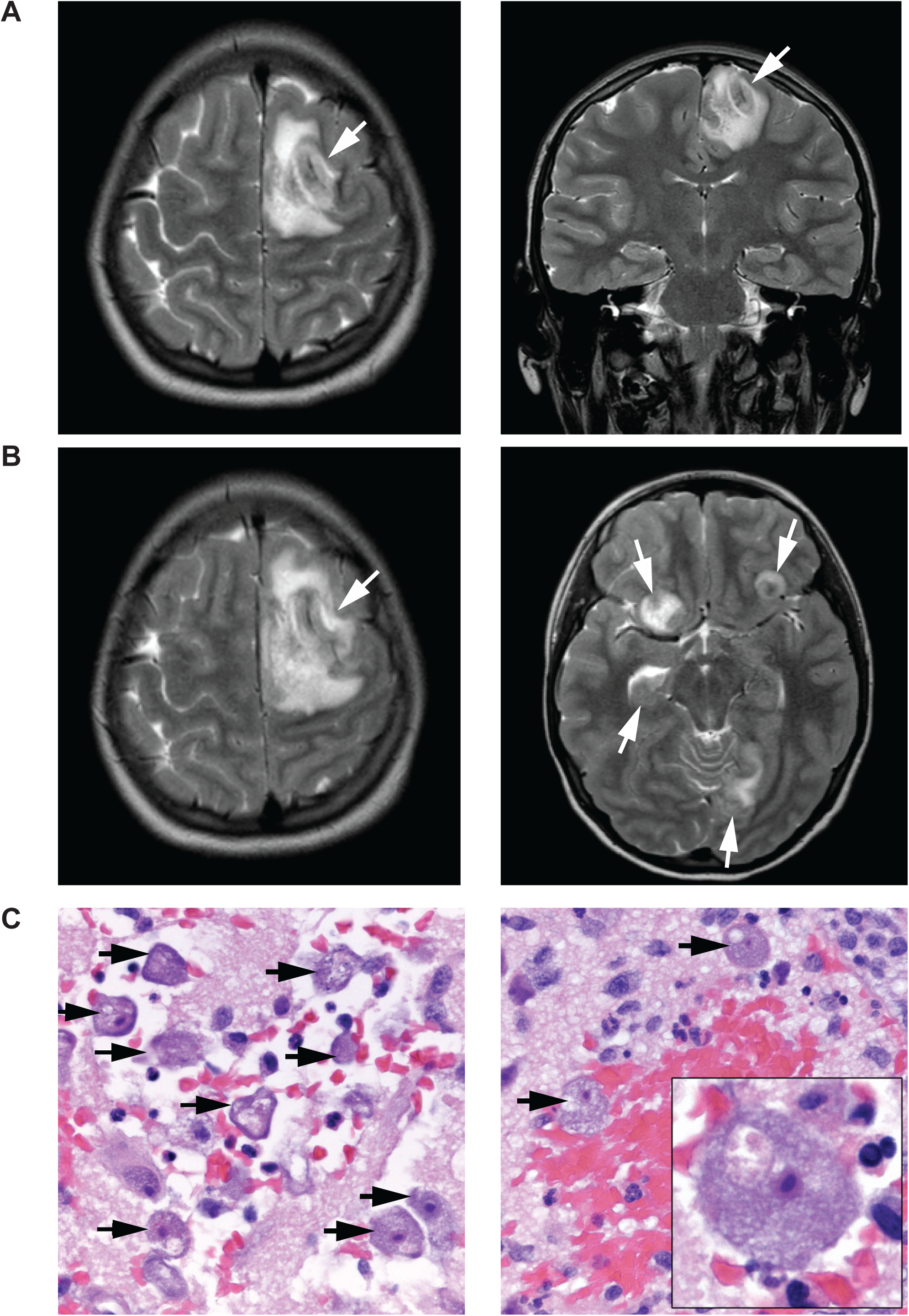
Magnetic resonance imaging and histopathology from a 15-year old patient with a fulminant acute encephalitis. **(A)** A hospital day (HD) 1 coronal T2-weighted MR image, demonstrating a hemorrhagic lesion with surrounding edema within the superior left frontal lobe (left panel, white arrow) and left occipital lobe (right panel, white arrow). **(B)** A HD 5 contrast-enhanced T1 weighted MR image, revealing enlargement of the pre-existing left frontal lobe lesion (left panel, white arrow), as well as interval development of numerous additional rim-enhancing lesions in multiple regions (right panel, white arrows). **(C)** 20X (left and right panels) and 100X fields of view (right panel, inset) of a brain biopsy specimen from the patient demonstrating numerous viable, large ameba (black arrows), with abundant basophilic vacuolated cytoplasm, round central nucleus, and prominent nucleolus, consistent with *Balamuthia mandrillaris.* There were areas of extensive hemorrhagic necrosis accompanied by a polymorphous inflammatory cell infiltrate including neutrophils and eosinophils (right panel).

Given the patient’s autoimmune predisposition and hemorrhagic appearance of the brain lesions, acute hemorrhagic leukoencephalitis was initially suspected and intravenous methylprenisolone (1,000 mg daily) was given HD 2-5. The patient clinically deteriorated with worsening headache, increasing weakness, and altered mental status on HD 5. Repeat MRI on HD 5 demonstrated enlargement of the previous hemorrhagic lesions with interval development of multiple rim-enhancing lesions (Fig. 4B). Steroids were discontinued and broad-spectrum antimicrobial therapy with vancomycin, cefotaxime, metronidazole, amphotericin B, voriconazole and acyclovir was initiated. On HD 6, she underwent craniotomy for brain biopsy, revealing partially necrotic white matter, and had an external ventricular drain placed. CSF wet mount and gram stain, bacterial and fungal cultures, PCR testing for HSV and varicella-zoster virus (VZV), and oligoclonal bands were negative. Pathology of the brain biopsy sample showed a hemorrhagic necrotizing process with neutrophils, tissue necrosis, vasculitis and numerous amoebae. Parallel metagenomic NGS testing of CSF and brain biopsy samples confirmed the presence of sequences from *Balamuthia mandrillaris* (see below). She was additionally started on azithromycin, sulfadizine, pentamidine, and flucytosine on HD 7. On HD 8, she developed intracranial hypertension, cardiac arrest and died. Miltefosine had been requested and was en route from the CDC (Schuster et al. 2006; Martinez et al. 2010; Centers for Disease and Prevention 2013); however this medication did not arrive in time to administer before the patient died. Autopsy was not performed according to the wishes of the family.

### Identification of *Balamuthia* in CSF and brain biopsy material

Metagenomic NGS and SURPI bioinformatics analysis were used to analyze the patient’s HD 6 CSF and brain biopsy for potential pathogens. Analysis of the viral portion of RNA or DNA derived reads revealed only phages or misannotated sequences (Table S1), while most of the bacterial reads mapped to common skin / environmental contaminants such as *Propionibacterium* (12,926 reads) and *Staphylococcaceae* (6,028 reads) in the RNA library. In contrast, 79% (20,145 of 25,631) of non-chordate (lacking a backbone) eukaryotic reads that were taxonomically assigned at a species level from the RNA library were assigned to available 16S and 18S sequences of *Balamuthia mandrillaris* in the NCBI nt reference database (Booton et al. 2003) (Table S1; Fig. 5A). A minority of the non-chordate eukaryotic reads aligned to *Acathamoeba spp.* (145 reads). Reads to *Balamuthia* were also detected in the DNA (13 reads) and brain biopsy RNA libraries (8 reads). The coverage of the 16S rRNA gene in the RNA library was sufficiently high to assemble a 1,405 bp full-length contig sharing 99.9% identity with the 2046 strain of *Balamuthia*. In the 18S locus, mapped NGS reads from the patient spanned 98.1% of the gene and were 99.1% identical by nucleotide. No NGS hits were detected to the RNAseP gene, the only additional *Balamuthia* gene represented in the NCBI nt reference database as of August 2015.

**Figure 5.**
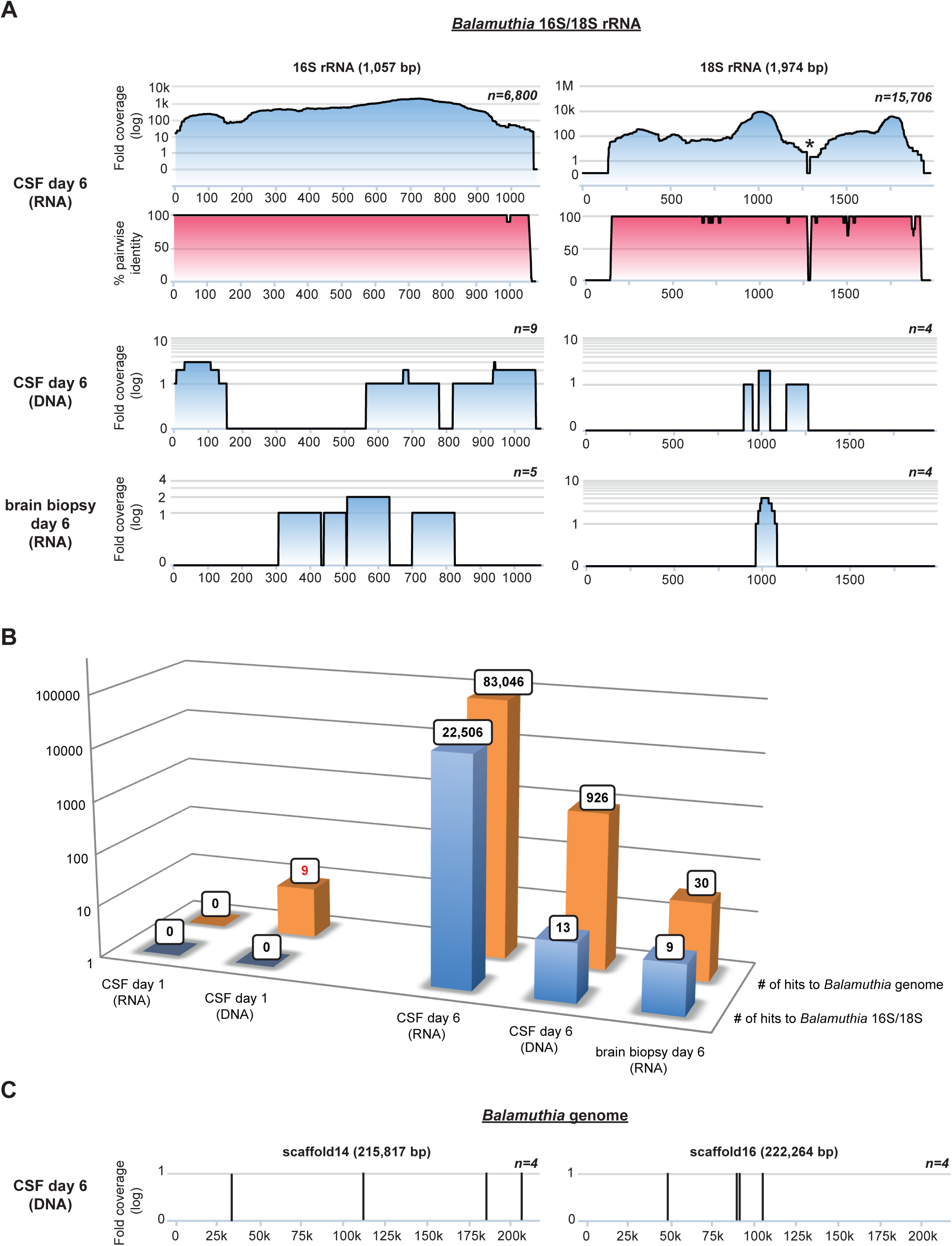
Identification of *Balamuthia mandrillaris* infection by metagenomic next-generation sequencing (NGS). **(A)** Coverage maps (blue gradient) and pairwise identity plots (magenta gradient) of 2 of the 3 available sequences from *Balamuthia* (16S/18S rRNA genes) in the NCBI nt reference database as of August 2015 and prior to sequencing of the draft genome. Shown are coverage maps corresponding to day 6 DNA and RNA libraries from CSF and a day 6 mRNA library from brain biopsy. No hits to 16S and 18S *Balamuthia* sequences were seen from day 1 samples. The asterisk denotes an area with artificially low coverage after taxonomic classification of the NGS reads due to high conservation among eukaryotic sequences (e.g. human, *Balamuthia*, etc.) within that region **(B)** A bar graph of the number of species-specific NGS reads aligning to *Balamuthia* 16S/18S rRNA (blue) or the *Balamuthia* genome (orange) in day 1 or day 6 samples. Note that with the availability of the newly assembled 44Mb *Balamuthia* genome, diagnosis of *Balamuthia mandrillaris* encephalitis at day 1 would have possible by detection of 9 species-specific reads (red boldface). **(C)** Coverage maps of two large scaffolds, ∼216 kB and ∼222 kB in size, from the *Balamuthia* draft genome, showing 8 out of 926 hits to *Balamuthia* in the day 6 CSF DNA library that are identified by SURPI after the draft genome sequence is added to the reference database (versus only 13 hits previously).

We then sought to determine in retrospect whether earlier detection and diagnosis of *Balamuthia* infection in the case patient by NGS would have been feasible. Metagenomic NGS of a day 1 CSF sample followed by SURPI analysis using the June 2014 NCBI nt reference database generated no sequence hits to *Balamuthia* (Fig. 5B; Table S1). However, repeating the analysis after adding the draft genome sequence of *Balamuthia mandrillaris* to the reference database resulted in the detection of many additional *Balamuthia* reads (Fig. 5B and C) Importantly, 9 species-specific DNA reads were detected from day 1 CSF (Fig. 5B, boldface text; Table 3). Although only 2 of 9 putative *Balamuthia* reads had identifiable translated nucleotide homology to any protein in the NCBI nr database, one of those reads was found to share 77% amino acid identity to the gluathione transferase protein from *Acanthamoeba castellani*, and hence most likely represented a *bona fide* hit to *Balamuthia*. These findings also indicated that the detection of *Balamuthia* reads was not due to errors in the draft genome assembly from incorporation of contaminating sequences from other organisms. Thus, detection of *Balamuthia* from the patient’s day 1 sample and a more timely diagnosis by metagenomic NGS would presumably not have been made without the availability of the full draft genome as part of the reference database used for alignment.

**Table 3.** *Balamuthia mandrillaris* reads from the day 1 patient sample.

| Read # | top NCBI NR match by BLASTx | % amino acid identity | E-value | matched scaffold in *Balamuthia* genome | % nucleotide identity (*Balamuthia* genome) | E-value (*Balamuthia* genome) |
| --- | --- | --- | --- | --- | --- | --- |
| 1 | none | – | – | scaffold46 | 96% | 5.00E-29 |
| 2 | none | – | – | scaffold26764 | 96% | 3.00E-27 |
| 3 | none | – | – | scaffold45 | 100% | 3.00E-36 |
| 4 | none | – | – | scaffold353 | 98% | 2.00E-24 |
| 5 | fumareylacetoacetase [Flavobacteriales bacterium BRH_c54] | 82% | 3.00E-05 | scaffold203 | 100% | 3.00E-36 |
| 6 | none | – | – | scaffold106 | 96% | 1.00E-26 |
| 7 | glutathione transferase protein [Acanthamoeba castellani str. Neff] | 77% | 0.022 | scaffold30 | 99% | 8.00E-34 |
| 8 | none | – | – | scaffold112 | 97% | 7.00E-28 |
| 9 | none | – | – | scaffold511 | 98% | 5.00E-26 |

## DISCUSSION

In this study, we describe a “virtuous cycle” of clinical sequencing in which the continually increasing breadth of microbial sequences in reference databases improves the sensitivity and accuracy of infectious disease diagnosis, in turn driving the sequencing of additional reference strains. The assembly of the first draft reference genome for *Balamuthia* not only enhances the potential sensitivity of metagenomic NGS for detecting this pathogen, as shown here, but also provides target sequences such as the rps3 intron / intergenic region or high-copy number 28S rRNA gene that can be leveraged for the future development of more sensitive and specific diagnostic assays. Given the lack of proven efficacious treatments for *Balamuthia* encephalitis, it is unclear whether even a much earlier diagnosis at HD 1 would have impacted the fulminant course of our case patient’s infection. However, it has been suggested that timely intervention in cases of *Balamuthia* might lead to improved outcome (Bakardjiev et al. 2003). In addition, promising new experimental treatments such as miltefosine (Schuster et al. 2006; Martinez et al. 2010; Centers for Disease and Prevention 2013), administered to the survivor infected by the sequenced 2046 strain (Vollmer and Glaser, submitted), are now available.

Unbiased metagenomic NGS is a powerful approach for diagnosis of infectious disease because it does not rely on the use of targeted primers and probes, but rather, detects any and all pathogens on the basis of uniquely identifying sequence information (Chiu 2013). Rapid and accurate bioinformatics algorithms (Zaharia et al. 2011; Wood and Salzberg 2014; Buchfink et al. 2015; Freitas et al. 2015) and computational pipelines (Naccache et al. 2014) have also been developed, with the capacity to analyze metagenomic NGS data in clinically actionable time frames. Nevertheless, we demonstrate here the critical role of comprehensive reference genomes in the NGS diagnostic paradigm. The availability of pathogen genomes with coverage of all clinically relevant genotypes can maximize the utility of NGS in not only diagnosis of individual patients (Wilson et al. 2014; Naccache et al. 2015), but also for public health applications such as transmission dynamics (Grad and Lipsitch 2014) and outbreak investigation (Briese et al. 2009; Greninger et al. 2015b).

In the field of amoebic encephalitis, draft genomes are now available for *Acanthamoeba castellani* (Burger et al. 1995), *Naegleria fowleri* (Zysset-Burri et al. 2014), and *Balamuthia* mandrillaris. However, more sequencing is certainly necessary to better understand the genetic diversity of these eukaryotic pathogens. In particular, shotgun sequencing and comparative analysis of mitochondrial genomes from 7 *Balamuthia* strains uncovered at least 3 unique lineages, one of which was comprised entirely of amoebae isolated from California, revealing that geographic differences likely exist among strains (Fig. 1B). This study also identified a unique locus in a putative rps3 intron/intergenic in the mitochondrial genome that is an attractive target for a clinical genotyping assay (Figs. 1C and 3). Given the rarity of the disease, it is unknown whether infection by different strains of *Balamuthia* would affect clinical course or outcome, although the availability of routine genotyping could help in addressing this question.

Limitations to this study include the small number of accessible clinical samples of *Balamuthia mandrillaris* infection and assembly of a draft genome with >14,000 scaffolds as a result of restricting the sequencing to short reads. The use of long read technologies based on single molecular, real-time (SMRT) or nanopore sequencing will likely be needed to achieve a highly contiguous, haploid genome. Furthermore, additional RNA sequencing of *Balamuthia* will be needed to predict transcripts, identify splice junctions, and enable complete annotation of the genome.

In summary, we demonstrate here that the availability of pathogen reference genomes is critical for the sensitivity and success of unbiased metagenomic next-generation sequencing approaches in diagnosing infectious disease. In hindsight, more timely and potentially actionable diagnosis at hospital day 1 in a fatal case of PAM from *Balamuthia mandrillaris* would have required the availability of the full genome sequence. Thus, in addition to revealing a significant amount of evolutionary diversity, the draft genome of *Balamuthia mandrillaris* presented here will improve the sensitivity of sequencing-based efforts for diagnosis and surveillance, and can be used to guide the development of targeted assays for genotyping and detection. The draft genome also constitutes a valuable resource for future studies investigating the biology of this eukaryotic pathogen and its etiologic role in PAM.

## METHODS

### Ethics

Informed consent was obtained from the patient’s parents for analysis of her clinical samples. This study was approved by the Colorado Multiple Institutional Review Board (IRB). Coded samples were analyzed for pathogens by NGS under protocols approved by the University of California, San Francisco IRB.

### Metagenomic Sequencing of CSF and Brain Biopsy

Total nucleic acid was extracted from 200 μL of CSF using the Qiagen EZ1 Viral kit. Half of the nucleic acid from CSF was treated with Turbo DNase (Ambion). Total RNA was extracted from 2 mm^3^ brain biopsy tissue using the Direct-zol RNA MiniPrep Kit (Zymo Research), followed by mRNA purification using the Oligotex mRNA Mini Kit (Qiagen). Total RNA from CSF and mRNA from brain biopsy was reverse-transcribed using random hexamers and randomly amplified as previously described (Greninger et al. 2015b). The resulting double-stranded cDNA or extracted DNA from CSF (the fraction not treated with Turbo DNase) was used as input into Nextera XT, following the manufacturer’s protocol except with reagent volumes cut in half for each step in the protocol. After 14-18 cycles of PCR amplfiication, barcoded libraries were cleaned using Ampure XP beads, quantitated on the BioAnalyzer (Agilent), and run on the Illumina MiSeq (1 × 160 bp run). Metagenomic NGS data were analyzed for pathogens via SURPI using NCBI nt/nr databases from June 2014 (Naccache et al. 2014).

A rapid taxonomic classification algorithm based on the lowest common ancestor was incorporated into SURPI, as previously described (Greninger et al. 2015b), and used to assign viral, bacterial, and non-chordate eukaryotic NGS reads to the species, genus, or family level. For the SNAP nucleotide aligner (Zaharia et al. 2011), an edit distance cutoff of 12 was used for viral reads (Naccache et al. 2014), but adjusted to a more stringent edit distance of 6 for bacterial and non-chordate eukaryotic reads to increase specificity.

### Propagation of Balamuthia mandrillaris in culture

Trypsin-treated cultures from Vero or BHK cell monolayers and *Balamuthia mandrillaris* 2046 strain amoebas were placed into T25 culture flasks in Dulbecco’s Modified Eagle Medium (DMEM) plus 10% bovine serum, 1% of the antibiotics 100 U/mL penicillin, 0.1 mg/mL streptomycin, and 0.25 μg/mL fungizone and incubated at 37° CO_2_ for 7 to 10 days plus 2 days at room temperature until the underlying cell sheet was completely destroyed and only actively dividing amebae were seen floating and/or attached to the surface. Attached amoebas were freed by gently tapping the side of the flask or putting the flask on a bed of ice for 20 minutes. The amoebas were concentrated by centrifugation for 5 minutes. After removal of supernatant, the amoeba pellet was washed by addition of PBS, centrifuged again, and then placed in a flask with Bacto-casitone axenic medium and allowed to grow for another 7-10 days, after which the amoebas were concentrated again, washed, and placed into fresh axenic medium. After a final centrifugation step, the amebae were collected, washed 3X in PBS, pelleted, and stored at -80° C.

The fluid drained, the ameba pellet was washed by adding PBS, centrifuged followed by placing the amoebas in a flask with Bacto-casitone axenic medium and allowed to grow for 7-10 days, collected again and placed into fresh axenic medium (Lares-Jimenez et al. 2015). The amebae were collected, washed 3X in PBS, pelleted, stored at -80°C.

### Sequencing and annotation of cultured *Balamuthia mandrillaris* 2046 strain

DNA from *Balamuthia mandrillaris* 2046 strain was extracted using the Qiagen EZ1 Tissue Kit and used as input for the Nextera Mate Pair Kit (Illumina) and Nextera XT Kit (Illumina), following the manufacturer’s instructions. Mate-pair libraries were sequenced on an Illumina MiSeq (2x80nt run and 2x300 nt), while the Nextera XT library was sequenced on an Illumina HiSeq (2x250bp paired-end sequencing) (Table 2). Mate-pair reads from run MP1 were adapter-trimmed with NxTrim (O’Connell et al. 2015), and the mitochondrial genome of strain 2046 and high-copy number contigs were assembled using SPAdes v3.5 (Bankevich et al. 2012; Greninger et al. 2015a). The average insert size of the mate-pair library was 2,187 nucleotides. Prediction of tRNA and rRNA genes were performed using tRNAscan-SE and RNAmmer v1.2, respectively (Lowe and Eddy 1997; Lagesen et al. 2007). ORFs were predicted in translation code 4 with the Glimmer gene predictor, and all predicted ORF sequences were confirmed using BLASTx and HHPred (Altschul et al. 1990; Soding et al. 2005).

Reads from runs MP2 and MP3 were mate-pair adapter-trimmed using NxTrim, while reads from all runs were quality-filtered (q30) and adapter-trimmed using cutadapt (Martin 2011). Reads that aligned to the *Balamuthia* mitochondrial genome and golden hamster (*Mesocricetus auratus*) were identified using SNAP (Zaharia et al. 2011) and removed prior to *de novo* assembly using platanus (Kajitani et al. 2014). Any scaffold of length less than 500 bp along with 62 scaffolds that aligned to *Mesocricetus auratus*, *Chlorocebus sabaeus*, Waddlia chondrophila, and *Enterobacteria phage phiX174* (all likely deriving from cell culture contamination), were removed.

### Quantitative RT-PCR

qRT-PCR of the *Balamuthia mandrillaris* 28S gene was performed using 20 μL total reactions of the Quantitect qRT-PCR Sybr Green kit (Qiagen) with 1 μL of extracted nucleic acid. Conditions were 50°C × 30 min, 95°C × 15 min, followed by 40 cycles of 95°C × 15 s, 60°C × 60 s using a final 0.5 μM concentration of each primer Bal-28S-F (5’-CTAGCCGTGCTGTAGAGTCG-3’) and Bal-28S-R (5’-CGGTCTCGAGCTTTTCCCTT-3’).

### Rps3 PCR confirmation

Genomic DNA was PCR-amplified using 0.5 μM final concentration of primers rps3-F (5’-CTGYTCGATTTTCGAAAAATAAAGTAG-3’) and rps3-R (5’-TGAAAGAAGAACATTTAGATCACGACT-3’) using 2X iProof HF Master Mix (Bio-Rad) in 20 μL total volume. Conditions were 95°C × 2 min, followed by 35 cycles of 95°C × 30 s, 52°C × 30 s, 72°C × 40 sec and a final incubation at 72°C × 2 min. PCR amplicons were visualized by 3% agarose gel electrophoresis.

## DATA ACCESS

The *Balamuthia mandrillaris* mitochondrial genomes have been deposited in NCBI under the following accession numbers: 2046 axenic (KP888565), 2046-1 (KT175740), V451 (KT030670), OK1 (KT030671), RP-5 (KT030672), SAM (KT030673), V188-axenic (KT175738), V188-frozen stock (KT175739), V039 (KT175741). The *Balamuthia mandrillaris* scaffolds have been deposited in NCBI WGS under the accession number LEOU00000000. Metagenomic NGS data from the brain biopsy and CSF fluid corresponding to non-human reads have been submitted to the NCBI Sequence Read Archive (SRA). NGS reads were filtered for exclusion of human sequences by both BLASTn alignment at an e-value cutoff of 10^-5^ and Bowtie2 high-sensitivity local alignment to the human hg38 reference database.

## ACKNOWLEDGEMENTS

This study was supported by grants from the National Institutes of Health (NIH) R01-HL105704 (to CYC) and UL1-TR001082 (to KM and SD), the Centers for Disease Control and Prevention (CDC) Emerging Infections Program U50/CCU915546-09 (CG), a University of California Discovery Award (CYC), and an Abbott Pathogen Discovery Award (CYC).

## AUTHOR CONTRIBUTIONS

ALG, KM, SD, and CYC conceived of and designed the study. ALG, TD, JB, SY performed the experiments. KM, DM, YN, MV, CAP, BKK, and SR took care of patients with *Balamuthia* infection and contributed clinical samples. KM, DM, CG, MV, CAP, BKK, SR, and CYC analyzed the clinical and epidemiological data. ALG, SNN, SF, JB, and CYC analyzed the genomic sequencing data. ALG, SNN, SF, and CYC developed and contributed software analysis tools. ALG, KM, and CYC wrote the manuscript.

---

**Table S1 – SURPI clinical metagenomic results**

